# Lifespan of male and female APP/PS1 and APP^NL-F/NL-F^ mouse models of Alzheimer’s disease

**DOI:** 10.1101/2024.10.15.618508

**Authors:** Hannah Roberts, Yimin Fang, Kathleen Quinn, Tiarra Hill, Mackenzie R. Peck, Andrzej Bartke, Kevin N. Hascup, Erin R. Hascup

**Author notes:** Erin R. Hascup, Department of Neurology, Dale and Deborah Smith Center for Alzheimer’s Research and Treatment, Southern Illinois University School of Medicine, P.O. Box 19628, Springfield, IL 62794-9628, USA.

## Abstract

Alzheimer’s disease (AD) disproportionately affects women, yet most preclinical research studies are male-centric. We performed lifespan analyses of male and female AD mouse models (APP/PS1 and APP^NL-F/NL-F^) and their shared genetic background control (C57BL/6). Survival curves support significant sex differences between within genotypes. Minimal longevity revealed increased age in male APP/PS1, and decreased age in APP^NL-F/NL-F^ mice. Maximal longevity revealed an increased average age in males. Furthermore, median lifespan differed between sex and genotype. This study supports sexual dimorphic survival in two mouse models of AD, emphasizing the need to examine mechanisms and treatments in both sexes.

## INTRODUCTION

Alzheimer’s disease (AD) leads to progressive loss of memory and cognition, with aging considered the predominant risk factor. Although women have a longer lifespan, their risk of developing AD and the incidence of the disease is higher compared to age-matched men [1]. Women account for two thirds of AD cases in the United States, experience greater cognitive deterioration, and have broader dementia-related behavioral symptoms [2]. Despite this, the majority of preclinical AD research has historically been conducted in male mice. While recent research has examined sex-based mechanistic differences, survival rates have not been determined [3]. To address potential sex-based differences in healthspan in AD, we examined survival in two AD mouse models (APP/PS1 and APP^NL-F/NL-F^) and their shared genetic background control (C57BL/6). Our laboratory has shown sex differences in cognition and metabolism [4-12]; however, little is known about their biological aging differences that could be a contributing factor when making chronological aging comparisons.

Double transgenic APP/PS1 mice overexpress amyloid precursor protein (APP) and presenilin 1 (PS1) genes. Disease onset and severity is accelerated compared to some knock-in mouse models [13]. APP/PS1 mice express elevated soluble amyloid-β (Aβ)_42_ levels by 2-4 months of age that progress into plaque pathology [14, 15]. Changes in neurotransmission have been reported as early as 2-4 months, while cognitive decline and a visible accumulation of insoluble Aβ_42_ are observed at approximately 6-8 months of age in both male and female mice, with significant cognitive impairment by 10-12 months [14, 16]. APP/PS1 mouse models have also shown sex-related differences with earlier pathology and increased β-secretase processing observed in females [6, 7, 17].

APP^NL-F/NL-F^ knock-in mice are a model of AD that express a humanized APP with both the Swedish and Iberian mutations. Differing from the APP/PS1 model, APP^NL-F/NL-F^ mice express APP at wild-type levels to avoid an overexpression of APP [13] and are lacking PS mutations [18]. APP^NL-F/NL-F^ mice have increased total Aβ_42_ production and eventual oligomerization and plaque formation characteristic of AD pathology [7]. Plaque deposition occurs in the APP^NL-F/NL-F^ strain for both sexes as early as 6 months with behavioral changes occurring by 18 months of age [7, 18]. Sex differences have also been observed in APP^NL-F/NL-F^, but few studies have reported on these changes [5, 7, 18, 19]. Interestingly, certain behavioral aspects including reversal learning and attention deficits, in female APP^NL-F/NL-F^ mice have no significant difference compared to males [18]. Male APP^NL-F/NL-F^ have shown impaired spatial long-term memory during Morris Water Maze (MWM) tests, but do not show loss in recognition memory [5]. Another study shows 12-month-old APP^NL-F/NL-F^ males performed significantly worse on the MWM task compared to age- and genotype-matched females [7].

Despite known sex differences in the human population with AD and recent insight into sexual dimorphism in mouse models of AD, little is known about the lifespan of the various mouse models of AD. In this study, we assessed sex differences in survival and longevity in APP/PS1 and APP^NL-F/NL-F^ mice to underscore the disparate healthspan in relation to known amyloid pathology. The information presented in this manuscript will aid with future experimental design and provide a clearer understanding of lifespan and biological aging across the sexes of these two AD models.

## MATERIALS AND METHODS

### ANIMALS

Protocols for animal use were approved by the *Institutional Animal Care and Use Committee* at Southern Illinois University School of Medicine (Protocol #2022-055). Parental strains of C57BL/6 (RRID:IMSR JAX:000,664) and APP/PS1 (RRID:MMRRC_034832-JAX) originated from The Jackson Laboratory (Bar Harbor, ME), while APP^NL-F/NL-F^ (RRID: IMSR_RBRC06343) founder breeders were obtained from Riken (Japan). Pregnant dams were housed individually with nesting material until after weaning pups. Post-weaning, mice were group housed based on sex and genotype on a 12:12 hr light:dark cycle, and food (LabDiet PMI Feeds Chow 5001) and water were available *ad libitum*. Genotypes were confirmed by tail tip DNA testing (TransnetYX, Inc; Cordova, TN). All mice were monitored from date of birth until date of natural death or at veterinarian’s request for moribund/health issues using TransnetYX Colony management software (Breeding Management Software; V1.0.205). Age was recorded by number of days alive. Mice with the following criteria were excluded: non-genotype strains, inconclusive or unknown genotype, experimental or breeding mice, unknown sex, unknown breeding origin, and those euthanized for reasons other than moribund issues at the request of the veterinarian.

### DATA ANALYSIS

We incorporated a retrospective study design whereby data was retrieved from colony maintenance software (TransnetYX Colony). Kaplan-Meier survival distributions of percent survival probability with 95% confidence intervals were constructed based on number of days alive utilizing log-rank (Mantel-Cox) to assess significant differences between genotypes and sexes. Minimal and maximal longevity were calculated by averaging age across the youngest and oldest 20% survived, respectively. Averages of minimal and maximal longevity were compared using independent t-tests to assess significant age differences between genotypes and using one-way ANOVA (Tukey’s multiple comparisons) to analyze significant age differences between sexes. Median and interquartile range of survival times show survival days to 75, 50, and 25% of mice remaining for each respective genotype.

Significance for all tests were determined at p < 0.05. All data are presented as mean ± SEM and statistical analyses and figure construction were performed with GraphPad Prism (GraphPad Prism 9 Software, Inc., La Jolla, CA; RRID:SCR_002798).

## RESULTS

Survival analyses indicate significantly shorter lifespan for females compared to males for the C57BL/6 and APP/PS1 genotypes (C57BL/6: χ^2^ = 9.38, df = 1, *p* < 0.0001; APP/PS1: χ^2^ = 161.1, df = 1, *p* < 0.0001; Fig. 1A). APP^NL-F/NL-F^ males and females have similar rates of survival (APP^NL-F/NL-F^: χ^2^ = 8.69, df = 1, *p* < 0.0001; Fig. 1A). Additionally, female APP/PS1 mice had the shortest survival rate compared to C57BL/6 and APP^NL-F/NL-F^ mice, whereas female APP^NL-F/NL-F^ mice had the longest survival rate (χ^2^ = 322.6, df = 2, *p* < 0.0001). C57BL/6 mice had the longest survival rate for males between the genotypes, whereas APP^NL-F/NL-F^ mice had the shortest survival rate (χ^2^ = 43.66, df = 2, *p* < 0.0001) (Fig. 1B).

**Figure 1.**
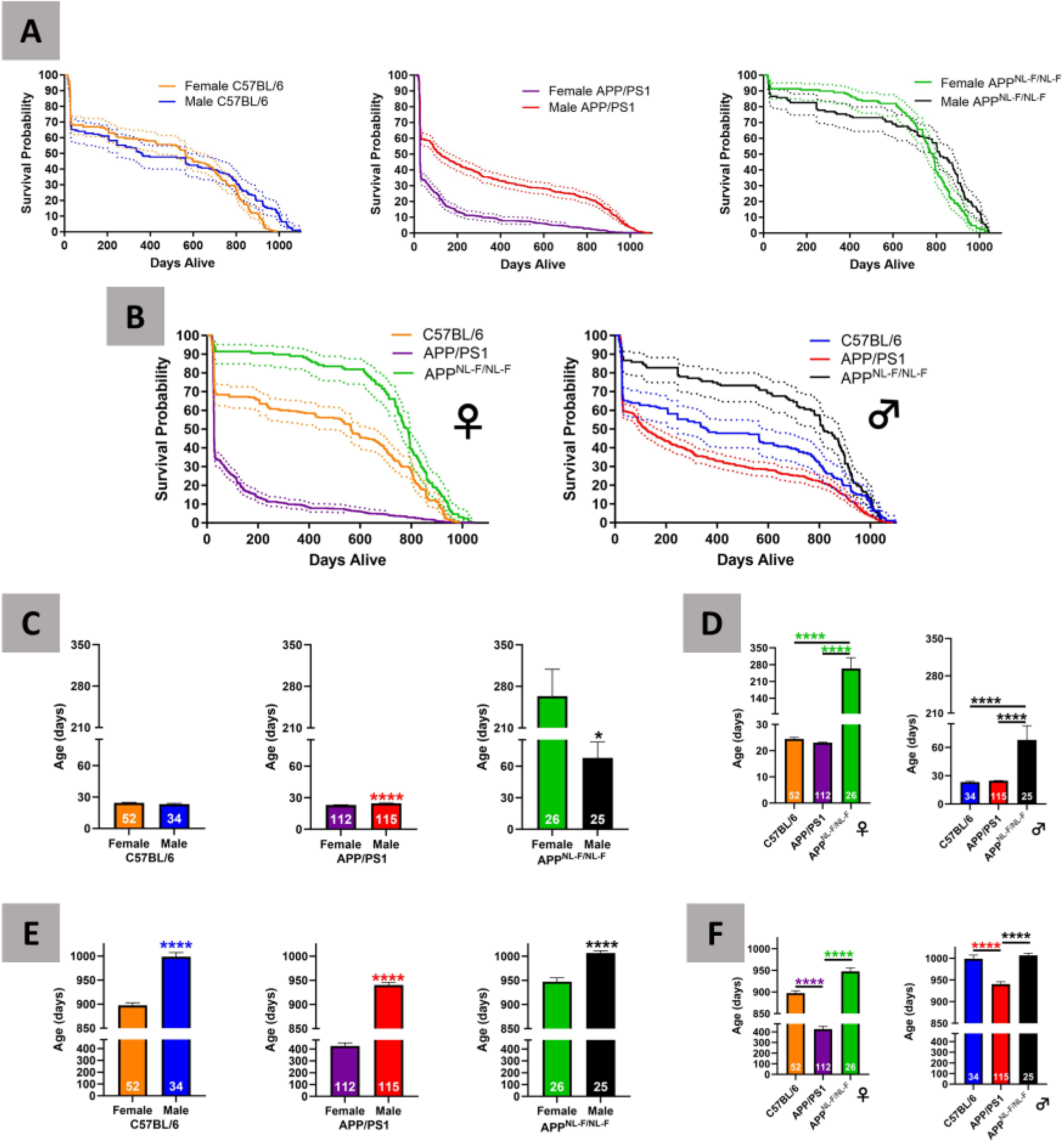
Survival and longevity analysis of C57BL/6, APP/PS1, and APP^NL-F/NL-F^ mice. Kaplan Meier curve is shown as a solid line, 95% confidence limits are expressed as dotted lines. Survival curves comparing between sex (Fig. 1A) and between genotypes (Fig. 1B). Minimal longevity between sex (Fig. 1C) and genotype (Fig. 1D) comparisons for the shortest surviving 20%. Maximal longevity between sex (Fig. 1E) and genotype (Fig. 1F) comparisons for the oldest surviving 20%.

Female (n = 52, mean: 24.50 ± 0.55) and male (n = 34, mean: 23.26 ± 0.74) C57BL/6 mice had comparable minimal longevity (C57BL/6 *t* = 1.4, df = 84, *p* = 0.18; Fig. 1C). Male APP/PS1 mice (n = 115, mean: 24.67 ± 0.19) had significantly longer minimal longevity than genotype-matched females (n = 112, mean: 23.06 ± 0.19; APP/PS1 *t* = 5.9, df = 225, *p* < 0.0001; Fig. 1C). Female APP^NL-F/NL-F^ mice (n = 26, mean: 263.70 ± 45.06) had significantly longer minimal longevity than males (n = 25, mean: 67.84 ± 15.06) of the same genotype (APP^NL-F/NL-F^ *t* = 4.1, df = 49, *p* = 0.0002; Fig. 1C). Female and male APP^NL-F/NL-F^ mice had the longest minimal longevity compared to sex-matched C57BL/6 or APP/PS1 mice [Females: F(2,187) = 91.6, *p* < 0.0001; Males: F(2,171) = 25.3, *p* < 0.0001; Fig. 1D], with no significant difference between same sex C57BL/6 and APP/PS1 mice.

Sex differences consisted of significantly shorter maximal longevity in female mice than male mice for all genotypes (C57BL/6: n = 52 and 34, mean: 897.4 ± 5.67 and 998.6 ± 8.55 respectively, *t* = 10.3, df = 84, *p* < 0.0001; APP/PS1: n = 112 and 115, mean: 425.50 ± 25.82 and 940.30 ± 5.53 respectively, *t* = 19.7, df = 225, *p* < 0.0001; APP^NL-F/NL-F^: n = 26 and 25, mean: 947.30 ± 7.97 and 1006 ± 4.98 respectively, *t* = 6.2, df = 49, *p* < 0.0001; Fig. 1E). Female and male APP/PS1 mice demonstrated the shortest maximal longevity among the three genotypes [Males: F(2, 171) = 25.6, *p* < 0.0001; Females: F(2,187) = 122.4, *p* < 0.0001; Fig. 1F]. There was no significant difference between either male or female C57BL/6 and APP^NL-F/NL-F^ for maximal longevity (Fig. 1F).

C57BL/6 female mice show a higher median survival time at 564 days compared to males at 359 days (Table 1). Sex-related differences were dichotomous for APP/PS1 mice as males demonstrate a median survival time of 120 days compared to females with 28 days (Table 1). APP^NL-F/NL-F^ male mice have a higher median survival time of 816 days compared to females at 782 days (Table 1).

**Table 1.** Median and interquartile range of survival times.

|  | Female |  |  | Male |  |  |
| --- | --- | --- | --- | --- | --- | --- |
|  | **C57BL/6** | **APP/PS1** | **APP<sup>NL-F/NL-F</sup>** | **C57BL/6** | **APP/PS1** | **APP<sup>NL-F/NL-F</sup>** |
| *N* | 257 | 562 | 128 | 172 | 573 | 126 |
| Survival Days to: |  |  |  |  |  |  |
| 75% Remaining | 28 | 25 | 655 | 28 | 27 | 339 |
| 50% Remaining | 564 | 28 | 782 | 359 | 120 | 816 |
| 25% Remaining | 805 | 104 | 859 | 850 | 694 | 927 |

## DISCUSSION

This study highlights sex differences in survival in mouse models of AD, which also supports differences in healthspan. Female APP/PS1 mice exhibited shorter lifespan than genotype-matched males in this study. This may be due to APP/PS1 female mice having higher soluble Aβ_40_ and Aβ_42_ by 4 months of age that progresses to greater plaque burden later in life than their age-matched male littermates [21] as well as increased microglial activation [22].

Female APP^NL-F/NL-F^ mice had a higher minimal longevity but a lower maximal longevity than genotype-matched males. In another study, APP^NL-F/NL-F^ female mice have shown a similar Aβ plaque burden to males at 2-5 months of age, however, female Aβ plaque burden was significantly increased at 12-16 months compared to males [23]. The observed sex-differences in this mouse strain could also be due to a disproportionate microglial response to Aβ in females, leading to faster disease progression [24]. The build-up of amyloid plaques due to an attenuated microglial clearance response may result in decreased rates of maximal survival for the APP^NL-F/NL-F^ female mice.

Genotype differences were present in APP/PS1 and APP^NL-F/NL-F^ mice compared to the C57BL/6 control. APP/PS1 mice feature an overexpression of wild-type APP, which can cause memory impairment with amyloid deposition also seen in APP^NL-F/NL-F^ mice [18]. Interestingly, APP^NL-F/NL-F^ female mice had the longest survival rate of the three genotypes assessed, possibly owing to this being a knock-in model with less severe pathology than APP/PS1. This is further supported by female APP/PS1 mice performing worse than APP^NL-F/NL-F^ mice in spatial learning and memory tasks [7].

Mice with higher survival past the minimal longevity point in this study could be more resilient to the detrimental effects and accumulation of AD pathology. Male mice have higher rates of maximal longevity in this study, which may suggest a resilience of males in AD pathology that is not present in females. Conversely, the more aggressive pathology observed in female AD mice may accelerate their biological aging thus shortening their lifespan compared to males [21]. Overall, our results show that female APP/PS1 mice have higher attrition rates and a sharper decline than males, which parallels proposed sex-based differences in disease progression and pathology in humans with AD [25]. Female APP^NL-F/NL-F^ mice show an initial sustained survival, but have a steeper, more dramatic decline in survival after midlife. Differences in survival in this group could be due to being a knock-in model with less severe disease pathology than transgenic strains [13].

Limitations for this study involve the reasoning behind sexually and genotypic dimorphism of survival rates. A possible explanation for the increased AD occurrence in elderly females as seen in older APP/PS1 female models is loss of reproductive hormones. Specifically, it has been shown that estrogen has neuroprotective effects in mitochondria exposed to Aβ [26]. The hippocampus, where AD pathology originates, has been shown to have high levels of estrogen receptors [23]. Estrogen decreases with age in both mice and humans [27, 28]. Female mice from the C57BL/6 genetic background control undergo reproductive senescence from 11 to 16 months of age [27]. As estrogen decreases with age in females, this could increase the incidence and hasten disease progression. Because males do not experience the same decrease in estrogen with age, this may explain why male AD mouse models in this study had greater maximal survival and why AD proportionately affects fewer men. Future studies will expand on the discrepancy in survival between sexes through further examination of estrogen present in female and male mice and how this decline may contribute to alterations in survival. Another limitation may be to explore the role of PS1 to understand why this leads to the least amount of survival for this group in males and females.

In conclusion, survival distributions indicated a significant difference between males and females for each genotype, and longer survival of males than females for the C57BL/6 and APP/PS1 groups. Same sex genotype comparisons support decreased survival distributions in APP/PS1 mice but increased in APP^NL-F/NL-F^ mice for both sexes. Previous male-centric AD preclinical research may have contributed to the high failure rate of clinical trials to show disease modifying benefits, underlining the importance of including both sexes in animal research to increase translatability of findings to the human population.

## Abbreviations: (alphabetically list all acronyms used)

(AD): Alzheimer’s disease
(APP): amyloid precursor protein
(PS1): amyloid-beta
(Aβ): presenilin 1

## DATA AVAIALBILITY

The data to support the findings of this study are available from the corresponding author upon reasonable request

